# Haplotype-aware multiomics unveils the regulatory basis of haplodiplontic life-cycle differentiation in a cosmopolitan marine alga

**DOI:** 10.1101/2024.05.26.595999

**Authors:** Tzu-Tong Kao, Ming-Wei Lai, Tzu-Haw Wang, Chia-Ling Yang, Miguel J. Frada, Chuan Ku

## Abstract

*Gephyrocapsa huxleyi* (formerly *Emiliania huxleyi*), a key coccolithophore alga influencing the global carbon cycle through photosynthesis and calcification, undergoes a haplodiplontic sexual life cycle with a calcifying non-flagellate diploid and a non-calcifying biflagellate haploid stage. To reveal the molecular basis of their morpho-physiological distinctions, we generated chromosome-level genome assemblies and compared the transcriptomes, proteomes, and methylomes for a pair of isogenic haploid and diploid model strains and conducted haplotype-aware analyses of their multiomic features. In addition to calcification and flagella, transcriptomes and proteomes of haploid and diploid cells modulate their differentiation in photosynthesis, sulfatases, DMSP degradation, DNA replication, and endomembrane system and transport. Haploid-diploid differential gene expression can be partially attributable to allelic imbalance (allele-specific expression) in diploid cells. Gene transcript abundance is positively associated with both CG and CHG gene-body DNA methylation, which can be inheritable, allele-specific, and differentiated between life-cycle phases. This multiomic study unravels the regulatory basis of unicellular algal life-cycle differentiation and provides valuable resources for investigating the ecologically important coccolithophore algae.

## Introduction

Sexual reproduction and transitions between life-cycle phases are hallmarks of eukaryotic evolution and a major mechanism for generating genomic variations in eukaryotes (Goodenough & Heitman, 2014). Sexual reproduction requires ploidy shifts between diploid and haploid genomes through meiosis and gamete fusion, of which the genetic toolkits date back to the last eukaryotic common ancestor (Sandra K. Floyd & John L. Bowman, 2007; Goodenough & Heitman, 2014). While animals typically have diplontic life cycles with haploid genomes restricted to gametes, land plants, various algae, and other eukaryotes are characterized by haplodiplontic life cycles, where both diploid and haploid cells can divide by mitosis, express distinct attributes, and form independent populations in microbial species (Umen & Coelho, 2019). The differentiation in morphological and physiological attributes between life-cycle phases in haplodiplontic life cycles expands the functional scope of these eukaryotes, where diploid and haploid organisms can explore different niches and display distinct life strategies (Frada *et al*., 2019; de Vries *et al*., 2021).

Coccolithophores are haplodiplontic, calcifying microalgae that are highly abundant and ecologically significant across global oceans (De Vargas *et al*., 2007; Taylor *et al*., 2017; Frada *et al*., 2019; de Vries *et al*., 2021). They contribute around 10% to global primary production as a major drive of the biological carbon pump (Rousseaux & Gregg, 2013). They also synthesize dimethylsulfoniopropionate (DMSP), a precursor of volatile dimethylsulfide, which may influence climate through cloud formation (Charlson *et al*., 1987; Alcolombri *et al*., 2015; Barak-Gavish *et al*., 2018). Most notably, coccolithophores fix carbon into CaCO_3_ through calcification, forming calcite plates (coccoliths) on the cell surface. Calcification not only releases CO_2_ inside the cell but contributes to carbon removal from the surface ocean with heavy coccoliths serving as ballast enhancing carbon export to the deep ocean (Taylor *et al*., 2017). The life cycle of coccolithophores alternates between a diploid and haploid phase with distinct morphological and ecological characteristics. *Gephyrocapsa huxleyi* (*sensu* Bendif *et al*., 2019); more commonly known as *Emiliania huxleyi*) is the most abundant and widespread species in the modern ocean and a model in coccolithophore research (Wheeler *et al*., 2023). Interestingly, for this species, coccolith formation is limited to the diploid phase. Diploid cells regularly produce heterococcoliths, while by contrast the haploid phase is not calcified and exclusively generates unmineralized organic scales that coat the cell surface and that are absent in the diploid cell (Fig. 1). In addition, diploid cells are non-motile and haploid cells biflagellate, diploid cells appear less sensitive to high light conditions (Houdan *et al*., 2005), and strikingly, haploid cells are resistant to specific lytic viruses that infect and kill diploid cells (Frada *et al*., 2008).

**Fig. 1.**
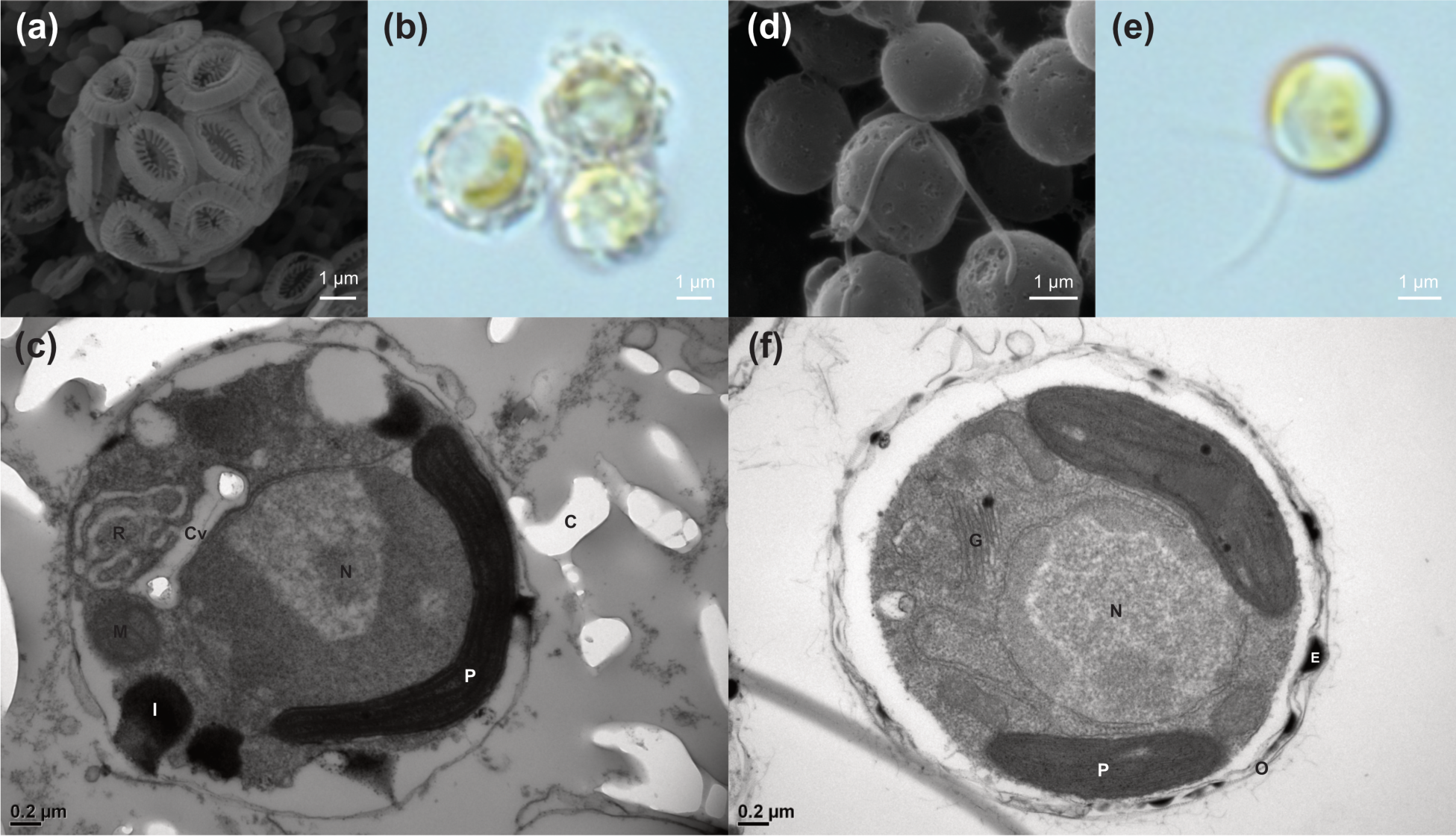
Morphology of *Gephyrocapsa huxleyi* RCC1216-G9 (diploid) and RCC1217-F3 (haploid). Scanning electron microscopy (a and d), differential interference contrast (b and e), and transmission electron microscopy (c and f) images of G9 and F3. C: empty space created by a fallen coccolith (calcite scale); Cv: coccolith vesicle; E: electron-dense particle; G: Golgi body; I: ion-dense vacuole; M: mitochondrion; N: nucleus; O: organic scale; P: plastid; R: reticular body.

Life cycle phase transition of *G. huxleyi* is poorly understood. A pair of diploid and haploid strains, RCC1216 and RCC1217, were identified and isolated from a diploid lab culture in 1999 (von Dassow *et al*., 2009) following a spontaneous meiosis event. A major limitation in past research has been the lack of a reference genome for the diploid-haploid pair. Previous genome sequences of *Gephyrocapsa* are derived from a diploid strain CCMP1516 (Read *et al*., 2013) or the closely diploid AWI1516 (Skeffington *et al*., 2023). Despite the very recent origin of *G. huxleyi* ∼0.3 million years ago (Bendif *et al*., 2019), its genomic structure displays substantial variability, with haploid genome size estimates ranging from 99 to 168 Mb and only ∼70% of genes shared across different strains (Read *et al*., 2013). Recent phylogenomic analyses further that RCC1216/RCC1217 and CCMP1516/AWI1516 belong to two distinct clades (Kao *et al*., 2022) or even species (*G. huxleyi sensu stricto* and *G. pseudohuxleyi*, respectively) (Bendif *et al*., 2023) with distinct morphology, biogeographic range, and plastid genomes (Kao *et al*., 2022). To better understand coccolithophore diploid-haploid differentiation at the molecular level, here we study RCC1216 and RCC1217 as a model system using a multiomics approach. Specifically, we aim to provide a comprehensive view of diploid-haploid differentiation at the molecular level and novel insight into its potential regulatory mechanisms.

## Materials and Methods

### Experimental Design

To investigate the life cycle phase differentiation of *G. huxleyi*, the model species of coccolithophores, we used a pair of diploid-haploid single-cell derived strains, RCC1216-G9 (G9) and RCC1217-F3 (F3). The strains RCC1216 and RCC1217 were originally obtained from the Roscoff Culture Collection (RCC) in 2012. To establish a robust foundation for downstream bioinformatic analyses, we constructed high-quality, haplotype-aware *de novo* genome assemblies for both strains. This involved combining data from long-read (PacBio and Oxford Nanopore) and linked-read (10x genomics plus Illumina) sequencing with chromosome optical mapping (Bionano) to identify existing genomic variations, including copy number variations, structural variations, insertion-deletions (Indels), and single-nucleotide polymorphisms (SNPs). For accurate transcriptome profiling, genome annotation, and gene expression analysis, we integrated data from Illumina RNA-Seq and full-length isoform sequencing (PacBio Iso-Seq). We also quantified and compared global protein expression using the DIA method. Lastly, we profiled genome DNA methylation, a known epigenetic regulatory mechanism for life cycle phase differentiation in some eukaryotes, by identifying 5-methylcytosine at symmetric (CG and CHG) and non-symmetric (CHH) sites using Illumina bisulfite sequencing and direct calling from kinetic data of the PacBio HiFi long reads. By integrating the omics data from these four components, our study provides a comprehensive view of diploid-haploid differentiation at the molecular level.

### Cell culture preparation

Single-cell derived cultures were generated using a Beckman Coulter MoFlo XDP cell sorter and cultured side-by-side under controlled conditions of ∼150 μmol light (16/8 hours light/dark), 18°C, and 60% relative humidity in a K/2 medium supplemented with an antibiotic cocktail of cefotaxime (50 ppm), carbenicillin (50 ppm), kanamycin (20 ppm), and amoxicillin (20 ppm). To maintain active growth, 1 ml of the cell culture was transferred into 40 ml of fresh medium every week. For DNA, RNA, and proteome extraction, the cells were harvested on the 6^th^ morning (at the late log growing phase) after the last refresh.

### Genome sequencing, assembly, and analyses

*DNA isolation and genome sequencing*—Approximately 1×10^8^ cells at the late log growing phase were harvested and embedded in a low-melting-point agarose plug for high-molecular-weight (HMW) DNA extraction. The cells were embedded in 0.75% Low-Melting-Point agarose (Lonza #50101, SeaPlaque^TM^) in SB buffer (25 mM Tris-HCl, 1 M D-sorbitol, and 25 mM EDTA) and digested with pH 9.5 DB buffer (250 mM EDTA, 1% *N*-lauroylsarcosine) containing 1 mg/ml proteinase K at 52°C for one day. The digested gels were subsequently washed with DB buffer (52°C, 30 mins, twice), digested with RNAase A (25µ1, 20 mg/mL) in DB buffer (1 hr, 37°C), stabilized with 1x Bio-Rad wash buffer, and DNA was retrieved by drop dialysis, following the Bionano Prep Cell Culture DNA Isolation protocol (#30026, Revision F). For PacBio and Nanopore sequencing, an additional step of salt:chloroform gDNA wash (https://www.pacb.com/wp-content/uploads/2015/09/Shared-Protocol-Guidelines-for-Using-a-Salt-Chloroform-Wash-to-Clean-Up-gDNA.pdf) was performed to remove polysaccharides. Genomic DNA was sequenced using PacBio (Sequel & Sequel IIe), Nanopore (MinION), 10x linked-read plus Illumina (NovaSeq 6000), or only Illumina (HiSeq 2500 and MiSeq) (Table S1). Additionally, chromosomes with labeled Nt.BspQI restriction sites (GCTCTTC) were optically mapped using Bionano (Saphyr).

*Genome assembly*—A hybrid approach of *de novo* genome assembly was used, integrating data from short-read, long-read, linked-read, and optical mapping. For the F3 genome, draft contigs were assembled with PacBio circular consensus sequences (CCS) using NextDenovo v2.5.0 (Hu *et al*., 2023). These contigs were then polished with PacBio HiFi and Illumina reads using NextPolish 1.4.0 (Hu *et al*., 2020), and non -eukaryotic contigs (Table S2) were removed by blasting against NCBI non-redundant nucleotide collection (nt; downloaded on Feb/19/2022). Eukaryotic contigs were scaffolded with 10x linked reads using ARCS (Yeo *et al*., 2018) and with nanopore reads using ARKS-long (Coombe *et al*., 2018). The genome was further hybrid-scaffolded with Bionano *de novo* genome maps using *hybridScaffold.pl* in Bionano Solve v3.6.1 and gap-filled with corrected long reads using TGS-GapCloser v1.1.1 (Xu *et al*., 2020). To validate the assembly correctness and copy number, the Illumina reads, PacBio reads, Nanopore reads/NextDenovo draft contigs, and Bionano molecules were mapped back to the superscaffolds. The bridged gaps were manually inspected, and only one (SS06 from coordinate 4,826k to 5,035k) lacked support from any long reads. The haplotype-aware phasing blocks were retrieved by calling haplotype-based variants using Freebayes v0.9.21 (Garrison & Marth, 2012) with short reads and phasing the variants with long reads using WhatsHap v0.19 (Patterson *et al*., 2015) In cases of partially duplicated superscaffolds (e.g. SS24 and SS26 of F3), haplotype-tagged reads were extracted to reconstruct *de novo* assemblies using NextDenovo and NextPolish.

*Genomic comparisons*—Genomic structural variations between the two genomes were analyzed by mapping the Bionano molecules of one strain to the genome of the other strain using *RefAligner* in Bionano Solve, and by aligning genome assemblies to each other using Minimap2 v2.15 (Li, 2018). Single nucleotide polymorphisms (SNPs) and short insertion-deletions (indels) were identified with GATK v4.3.0.0 (McKenna *et al*., 2010) that the variable sites were called using *HaplotypeCaller* and underwent hard filtering based on the recommended GATK *VariantFiltration* settings (SNPs: QD < 2.0, QUAL < 30.0, SOR > 3.0, FS > 60.0, MQ < 40.0, MQRankSum < -12.5, and ReadPosRankSum < -8.0; indels: QD < 2.0, QUAL < 30.0, FS > 200.0, and ReadPosRankSum < -20.0). The draft of variable sites was then used in base quality score recalibration using *BaseRecalibrator* and *ApplyBQSR* for subsequent variation calling. Convolutional neural net filtering was applied, and variants were annotated using GATK *HaplotypeCaller*, *CNNScoreVariants*, and *FilterVariantTranches*. The G9 variable sites were phased with long reads using WhatsHap. The synonymous and non-synonymous variable sites of each coding gene within and between G9 and F3 were analyzed using SNPGenie v2019.10.31 (Nelson *et al*., 2015).

### RNA sequencing, genome annotation, and functional annotation

*RNA sequencing*—Total RNA from G9 and F3 was extracted using the RNeasy Plant Mini Kit (Qiagen, Hilden, Germany). Each strain had four short-read libraries prepared using the TruSeq Stranded mRNA Library Prep Kit (Illumina, San Diego, CA, USA), following the manufacturer’s recommendations, and sequenced on Illumina NovaSeq 6000 in 150 bp paired-end mode. Additionally, one PacBio Iso-seq library was prepared using the SMRTbell® prep kit (PacBio, Menlo Park, CA, United States) and sequenced on the Sequel IIe.

*Genome annotation*—Repetitive sequences in the genome were modeled using RepeatModeler2 v2.0.2a (Flynn *et al*., 2020) and annotated with Dfam 3.5 and RepBase RepeatMasker Edition (20181026) using RepeatMasker v4.1.2 (Smit *et al*., 2015). Telomeric repeats were explored and annotated with the Telomere Identification toolKit (tidk) v0.2.0 (Tree of Life Kit, 2022) Both PacBio Iso-seq and Illumina RNA-seq reads were used for annotating the genome, which was soft-masked with repeats. PacBio Iso-seq CCS reads were mapped to the genome and then refined, clustered, and collapsed using PacificBiosciences Isoseq3 v3.8.2, with protein-coding regions predicted by GeneMarkS-T v.5.1 (Tang *et al*., 2015). The Illumina RNA-seq reads were aligned to the genomes and gene models were trained and predicted using GeneMark-ET and Augustus v3.4.0 in BRAKER 2.1.6 (Hoff *et al*., 2016). The two annotations were combined and renamed using TSEBRA v1.0.3 (Gabriel *et al*., 2021) for further isoform analyses, transcripts were filtered based on the Iso-Seq data using PacificBiosciences Pigeon. The shared genes and ortholog groups among genomes were analyzed using the amino acid datasets of haptophytes provided in the associated files of Skeffington *et al*. (2023) (DOI: 10.6084/m9.figshare.20463900) with OrthoFinder v2.5.2 (Emms & Kelly, 2019). Non-coding RNAs were predicted and annotated with the Rfam database v14.8 using structRNAfinder v1.0 (Arias-Carrasco *et al*., 2018).

*Functional annotation*—The functions of the predicted protein-coding transcripts were annotated using the EnTAP v0.10.8 pipeline (Hart *et al*., 2020) by searching against NCBI non-redundant protein collection (nr), UniProt Swiss-Prot, and UniProt TrEMBL databases (all downloaded on May/18/2022) using DIAMOND v0.9.9, and Gene Ontologies (GO), KEGG pathway orthologous groups (KO), and protein domains (SMART/Pfam) were explored and annotated based on the eggNOG database (downloaded on May/18/2022).

### Global protein identification and methylomic sequencing

*Global protein identification*—Total proteins were extracted and digested from three replicates of frozen cell pellets containing approximately 1×10^7^ cells of each strain, prepared according to Yun et al. (2020). The protein pellets were resuspended, filtered, and digested using the S-Trap-IMAC protocol (Chen *et al*., 2023, 2024). Mass spectrometry (MS) analyses were conducted on a Thermo Orbitrap Fusion Lumos Tribrid Mass Spectrometer using DIA mode.

*Methylomic sequencing***—**To prepare for bisulfite sequencing, cells were lysed using a modified lysis buffer (Tris 0.1M, EDTA 0.05M, NaCl 0.1M, SDS 1%, N-laurylsarcosine 2%, and proteinase K 1mg/ml in pH 8.0; modified from Bendif *et al*. (2019), and then genomic DNA was isolated following the PacBio Salt:Chloroform gDNA Wash protocol. Two samples of each strain were bisulfite converted using the EZ DNA Methylation Gold Kit (ZymoGenetics, Seattle, WA, USA). The two libraries were prepared using the TruSeq DNA Methylation Kit (Illumina) and subsequently sequenced using HiSeq in 150 bp and 250 bp paired-end modes, respectively.

### Statistical Analysis

*Transcriptomic Analyses*—The expression of transcripts was quantified with Illumina RNA-seq reads using Salmon v1.9.0 (Patro *et al*., 2017), correcting for GC bias (--gcBias) and performing 30 Gibbs sampling runs (--numGibbsSamples 30). Differential expression of genes and transcripts, as well as differential transcript usage, were analyzed with inferential replicate counts (i.e., the Gibbs from Salmon), normalized by the scaledTPM method using the Swish method carried out with the R package *fishpond* v2.0.1. The global allelic imbalance was tested using the SEESAW method (Wu *et al*., 2023). Differentially expressed genes and allelic imbalance genes were identified with q-values ≤ 0.01, while G9- or F3-specific genes were those with q-values ≤ 0.01 and a mean scaledTPM < 5 in the strain with fewer mapped reads.

*Proteomic Analyses*—The quantification and normalization of MS data were performed using the MaxLFQ method as implanted in DIA-NN v1.8.1 (Demichev *et al*., 2020). Differential protein expression was analyzed based on MaxLFQ value and the number of precursors (used for variance estimation) using the R package *DEqMS* v1.12.1 (Zhu *et al*., 2020). Differentially expressed genes were identified with q-values ≤ 0.05, while G9- or F3-specific genes were characterized by q-values ≤ 0.05 and absence of peptide detection in two or three out of three replicated samples in the strain with fewer mapped reads.

*Methylomic Analyses*—The bisulfite sequencing reads and PacBio HiFi reads that retained kinetics were mapped to the genomes and used for methylation calling with BSBolt v1.4.5 (Farrell *et al*., 2021) and PacificBiosciences Primrose v1.3.0, respectively. Genome-wide DNA methylation profiling was conducted using R package BSseq v.1.30.0 (Hansen *et al*., 2012), while their association with RNA expression was analyzed using MethGet (Teng *et al*., 2020). Differentially methylated CG, CHG, and CHH sites, genes, and regions were analyzed using the R package *DSS* v.2.42.0 (Wu *et al*., 2013). Differentially methylated sites were detected with an FDR threshold of ≤ 0.05, while differential methylation of genes and promoter regions (2000 bp upstream of genes) was determined either by mean methylation level (FDR ≤ 0.05; Datasets S2) or by overlapping with differential methylation regions surpassing areaStats thresholds. In comparisons with RNA expression, an additional set of regions, most correlated with RNA expression (coordinates -350∼-50 and 50-450 relative to start codon in CG context; and -200∼300 and 500-1500 in CHG context), were also assessed for differential methylation.

*Functional enrichment analyses*—Gene and transcript sets of interest, including copy number variation, differential gene/protein expression, transcript usage, allelic imbalance, and methylation levels, were analyzed for enriched GO, KO, SMART, and Pfam terms using the function “*enricher*” in R package *clusterProfiler* v4.2.2 (Wu *et al*., 2021). Additionally, enriched GO terms were analyzed using R package *topGO* v2.46.0 (Alexa & Rahnenfuhrer, 2021) with algorisms “classic” or “weight01”. Enriched GO terms identified by *clusterProfiler* (p < 0.05) were clustered using the MCL-based overlap coefficient in GOMCL v0.0.1 (Wang *et al*., 2020).

## Results

### Genome assemblies and annotations

Our *de novo* genome assemblies exhibit substantially improved genome contiguity compared with other assemblies of *Gephyrocapsa* (Figure 2A). The genome assembly of the haploid strain F3 is 151 Mb, comprised of 26+1 superscaffolds (SS), where SS26 is a partial duplicate of SS24 and not present in the diploid G9. The *G. huxleyi* genomes display a high GC content (∼66%) and contain a significant proportion of repetitive sequences (∼36% after excluding the sequences overlapped with genes; Fig. 2b and Table 1). The telomeric repeat sequence TTAGGG, commonly found in animals and other eukaryotes (Fulnečková *et al*., 2013) was detected in some superscaffolds but not all. The coding genes annotated utilizing both PacBio Iso-Seq full-length transcripts and Illumina RNA-seq reads amount to 35,323 in F3, with 4,005 (11%) of them absent in the CCMP1516 assembly (RefSeq GCF_000372725.1) (Read *et al*., 2013) or its still calcifying variant AWI1516 (Skeffington *et al*., 2023) (Fig. 2c). A total of 827 non-coding RNA genes were identified, including 349 rRNA genes that are mostly located on SS02, SS03, SS05, and SS06 (Fig. 2d).

**Fig. 2.**
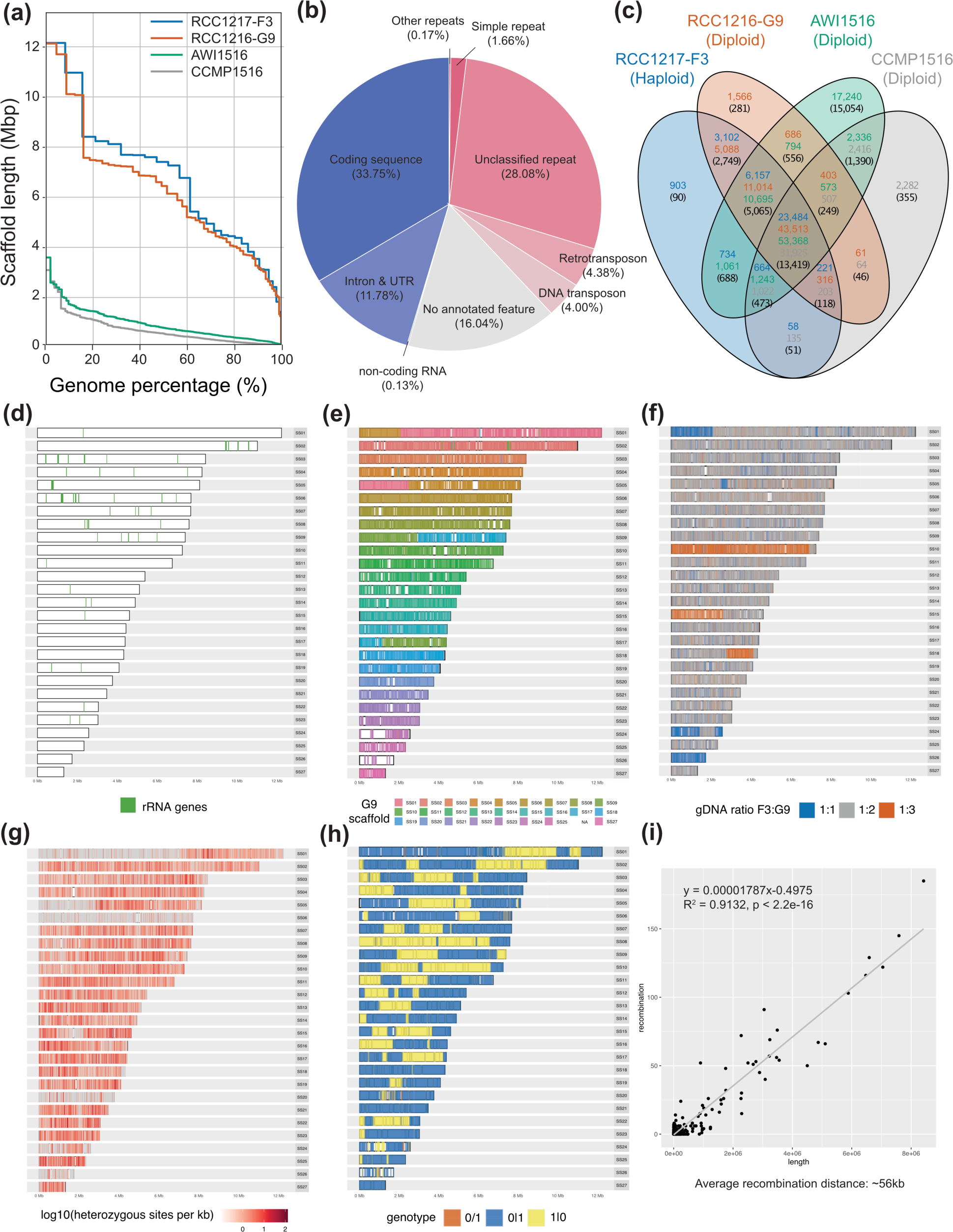
Genome assemblies, genomic features, structure variations, heterozygosity, and recombination rate. (a) N statistics (scaffold length at different percentages of genome assembly) of RCC1217-F3 and RCC1216-G9 (*G. huxleyi*) compared with CCMP1516 and AWI1516 (*G*. *pseudohuxleyi*). (b) Relative length percentage of coding and non-coding sequences of genes (blues) and different classes of repeats (reds) in F3. (c) Numbers of genes and orthogroups (in parenthesis) unique to or shared among the strains. (d) Distribution of rDNAs in F3. (e) Mapping of G9 superscaffolds to the F3 genome, revealing interchromosomal recombinations and large-scale indels. White: unmappable regions. (f) Distribution of gene copy number variations across the genome, showing partial aneuploidy. (g) Frequency of G9 heterozygous sites across the genome, with lower heterozygosity in certain genomic regions. (h) Genomic regions of F3 that have been phased into different G9 alleles (blue and yellow). Orange: un-phased regions in phasing blocks. White: regions outside of phasing blocks. (i) Correlation between the length of phasing blocks and the number of recombination sites.

**Table 1.**
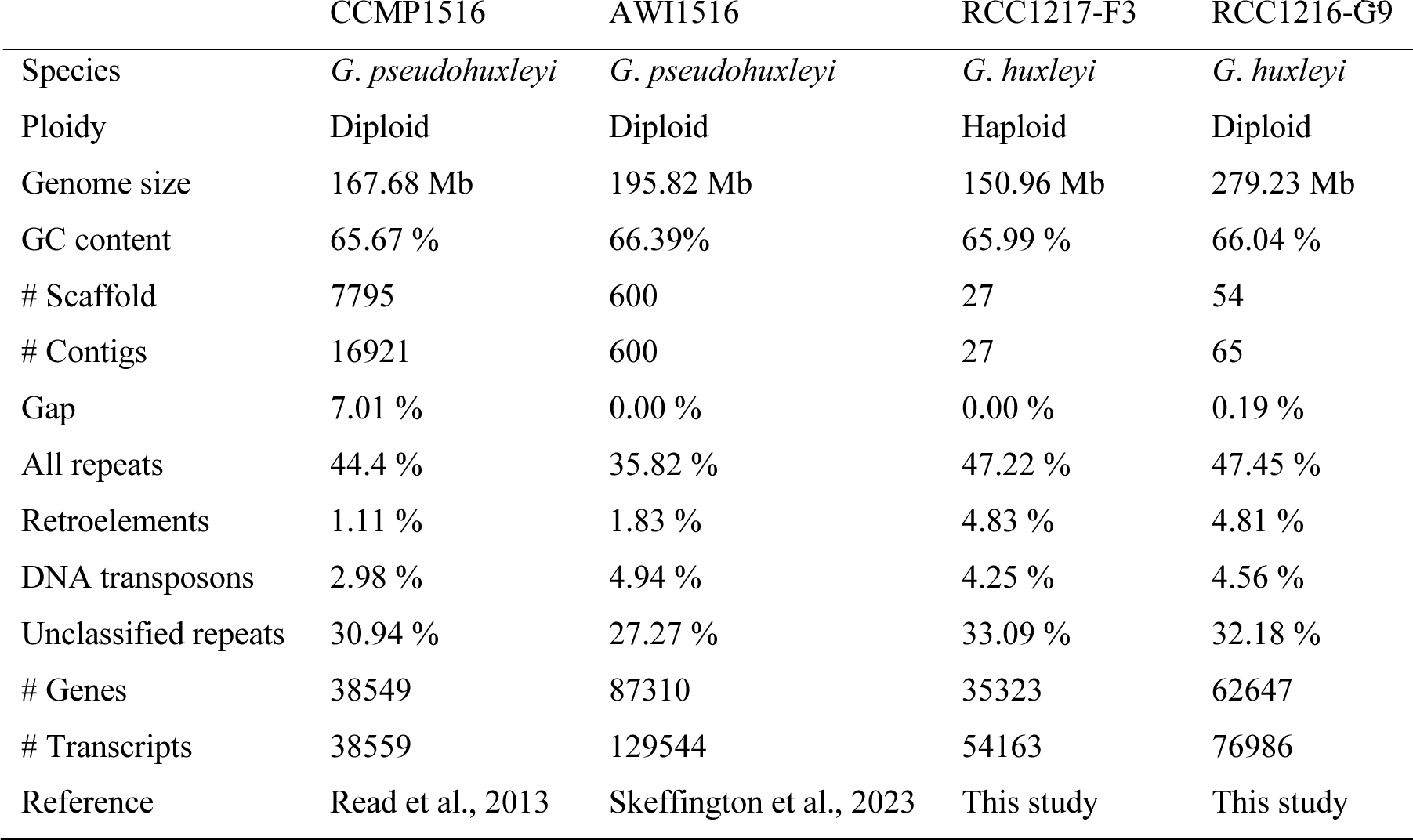
Statistics of the haploid and diploid genome assemblies of *Gephyrocapsa* spp.

### Genome variations and phasing

F3 and G9 genomes exhibit notable structural divergence (Fig. S1), including two chromosome-level recombination events between superscaffolds (SS01-SS05 and SS09-SS17; Fig. 2e), partial duplication of SS24 in F3, partial loss of SS01 in G9, and acquisition of partial copies of SS10, SS15, and SS18 in G9 (Fig. 2f). Noteworthy is a biased distribution of large indels (>350 bp), with a higher frequency of insertions in F3 (or deletions in G9) compared to F3 deletions (or G9 insertions), at a ratio of 444:97.

Approximately 0.33% (∼491 k) of sites within the G9 genome are heterozygotic, with the majority (99.87%) of these sites having one allele corresponding to the F3 haplotype. Specific regions in the G9 genome (e.g., SS06, SS20, and partial SS01 and SS05) display low heterozygosity (Figs. 2g, S2). Comparison with a single-cell-derived strain (B4) from another RCC1216 culture (kindly provided by the lab of Assaf Vardi) indicates that regions of low heterozygosity in G9 and B4 genomes mostly do not overlap, suggesting they are due to recent loss of heterozygosity after the separation of RCC1216 and RCC1217. A total of 13,405 sites (0.0089%) have diverged between F3 and G9, primarily concentrated within the regions of low heterozygosity, presumably due to the loss of the G9 allele corresponding to the F3 haplotype. We phased the variable sites with long reads, resulting in 213 phasing blocks covering 94.90% of the haploid genome (Fig. 2h; Dataset S1). The number of recombinations is positively correlated with the length of phasing blocks, with the average recombination distance estimated to be ∼56 kb (Fig. 2i).

By investigating the variation in gene copy numbers, as defined by gDNA ratios, of the two genomes, we found that only 75% of the genes possess two copies in the diploid genome relative to the haploid (Fig. 3a). Among the 25% that deviate from an expected 2:1 ratio, 12% have 3:1 ratio and 11% have a 1:1 ratio, which might be due to factors such as chromosome anomalies (e.g., aneuploidy) or gene duplications and losses. Notably, 2% of the genes have an even more pronounced deviation (with a ratio of < 0.5 or > 3.5 to 1) and were subsequently excluded from further analyses.

**Fig. 3.**
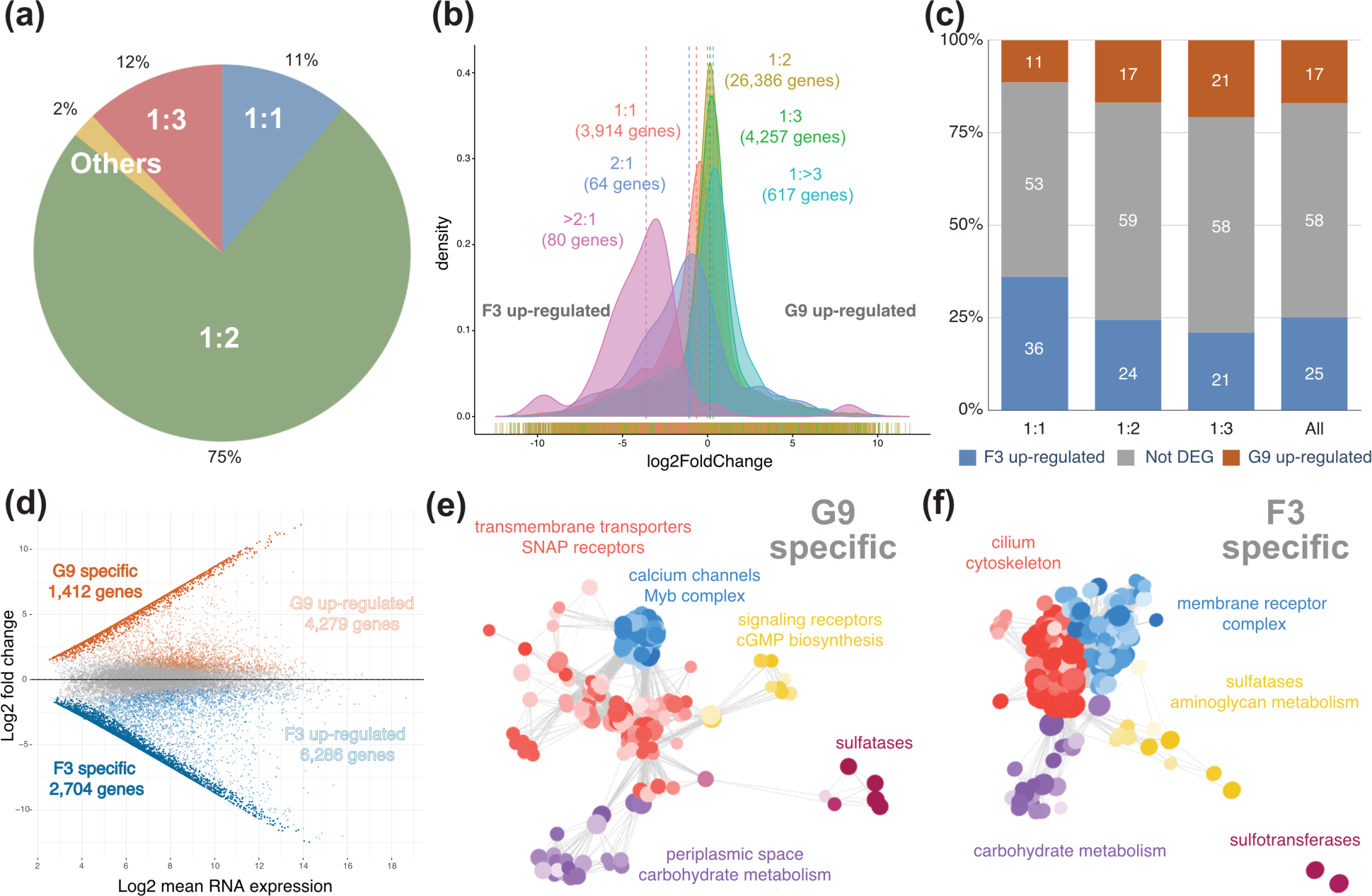
Gene copy number variation and transcriptomic differentiation. (a) Relative abundance of genes with different G9:F3 genomic DNA (gDNA) ratios. (b) Distribution patterns of RNA expression ratios for gene groups with different gDNA ratios. (c) Percentage of up- and down-regulated genes across different gDNA ratios. (d) MA plot of RNA expression, highlighting genes specifically expressed or up-regulated in each life cycle phases. (e) Clusters of enriched GO terms of diploid-specific transcripts. Each dot represents an enriched term, with darker colors indicating smaller p-values. Dots with similarity > 0.5 are linked by a grey line. Clusters with ≤ 5 genes were excluded. (f) Clusters of enriched GO terms of haploid-specific transcripts.

### Transcriptomic landscapes and gene functions

RNA differential expression comparisons between G9 and F3 were conducted, utilizing four replicates of Illumina RNA-seq samples (Table S1), collected on the sixth morning after transferring to a fresh 100 ml K/2 medium (starting with 100 cells/µl). Gene copy numbers correlate with RNA transcript abundance (Figs. 3b, 3c), but there is a high variance of RNA differential expression among genes with a specific haploid-diploid gDNA ratio (Fig. 3b), suggesting the impact of copy numbers on gene expression is minor. Out of the 25,224 genes with detectable transcripts, 1,412 genes (5.60%) have transcripts only in the diploid (mean scaled transcripts per million [scaledTPM] < 5 in F3) and 2,704 genes (10.72%) only in the haploid (mean scaledTPM < 5 in G9) (Fig. 3d). Given the phylogenetic distance of haptophytes from well-studied eukaryote models (animals, fungi, and plants), our understanding of haptophyte gene functions remains very limited. Among the transcribed genes, only 53.26% (13,434 genes) are associated with known functions, and the proportions for only diploid-expressed (33.92%; 479 genes) and only haploid-expressed (39.09%; 1,057 genes) genes are even lower (Dataset S2).

The functions enriched among genes only expressed in the diploid were divided into five clusters (Fig. 3e). The predominant cluster (red) primarily consists of membrane proteins, including genes implicated in coccolithogenesis, such as HCO_3_^−^ transporters in the solute carrier 4 family, cation and proton channels, antiporters, and vesicle fusion SNARE proteins. A closely related, highly compact cluster (blue) comprises Myb-like transcription factors and Ca^2+^ channels. The third cluster (purple) mainly encompasses genes involved in carbohydrate metabolism, including the glycoside hydrolase 3 family, beta-glucan synthase, and galactosyltransferase. Two smaller clusters, each containing around 20 genes, include guanylate cyclase receptors and adenylate guanylate cyclase (yellow) and sulfatases (magenta).

The predominant functional cluster (Fig. 3f; red) of the haploid-specifically expressed genes primarily consists of genes related to flagella assembly, intra-flagellar transportation, microtubule-based movement, nuclear division, and certain cell cycle regulatory proteins. Other clusters include membrane receptors (blue; ligand-gated channels, protein kinases, blue light photoreceptors, etc.), carbohydrate metabolism (purple; glycoside hydrolase 5 family, glycosyltransferase, cellulases, etc.), sulfatases plus aminoglycan-related glycosyltransferases (yellow), and sulfotransferases (magenta).

Using Pigeon in PacBio Transcript Toolkit, we identified isoforms in 6,092 genes, with 2,145 genes (35.21%) showing isoform differential expression patterns: 1,232 (20.22%) had up-regulated isoforms in the diploid, and 1,229 (20.17%) had up-regulated isoforms in the haploid. Genes with diploid up-regulated isoforms were enriched in plastid thylakoid membrane components, such as the light-harvesting complex, cytochrome b6f complex, and photosystem II genes (Dataset S3). Those genes with haploid up-regulated isoforms were enriched in mitotic cell cycle-related genes, such as cyclins, mitogen-activated protein kinases, and kinetochore complex components.

### Proteomic profiling

Global proteins were identified, quantified, and compared using three replicates of data-independent acquisition (DIA) data for G9 and F3 (Table S1). Due to limitations in protein detection, the number of genes with detectable proteins (7,116) is lower than that with RNA transcripts detected (25,224). There is a moderate correlation between protein and RNA expression (R = 0.52∼0.56; Figs. 3a-3c). Among genes with protein products detected, 1,082 are up-regulated in the diploid, with the majority (805 genes) being specifically detected in the diploid and enriched with transportation proteins (solute carriers, major facilitators, ion transporters, and antiporters) (Fig. 4e; Dataset S3). The 599 genes with protein expression up-regulated or only detected in the haploid are grouped into three enriched functional clusters: mini-chromosome maintenance (MCM) complex proteins, protein degradation enzymes (peptidases and proteasomes), and heat-shock proteins plus mitochondrial matrix proteins (dehydrogenases and reductases) (Fig. 4f). It is notable that flagella- and cell motion-related genes that were highly upregulated in the RNA differential expression profile were not enriched in the haploid proteome. Unlike RNA, which is typically localized within the main cell body, flagella are easily detached from the cell body during microscopic observations. Therefore, the absence of these proteins may be due to the loss of flagella during the pelleting step of sample preparation.

**Fig. 4.**
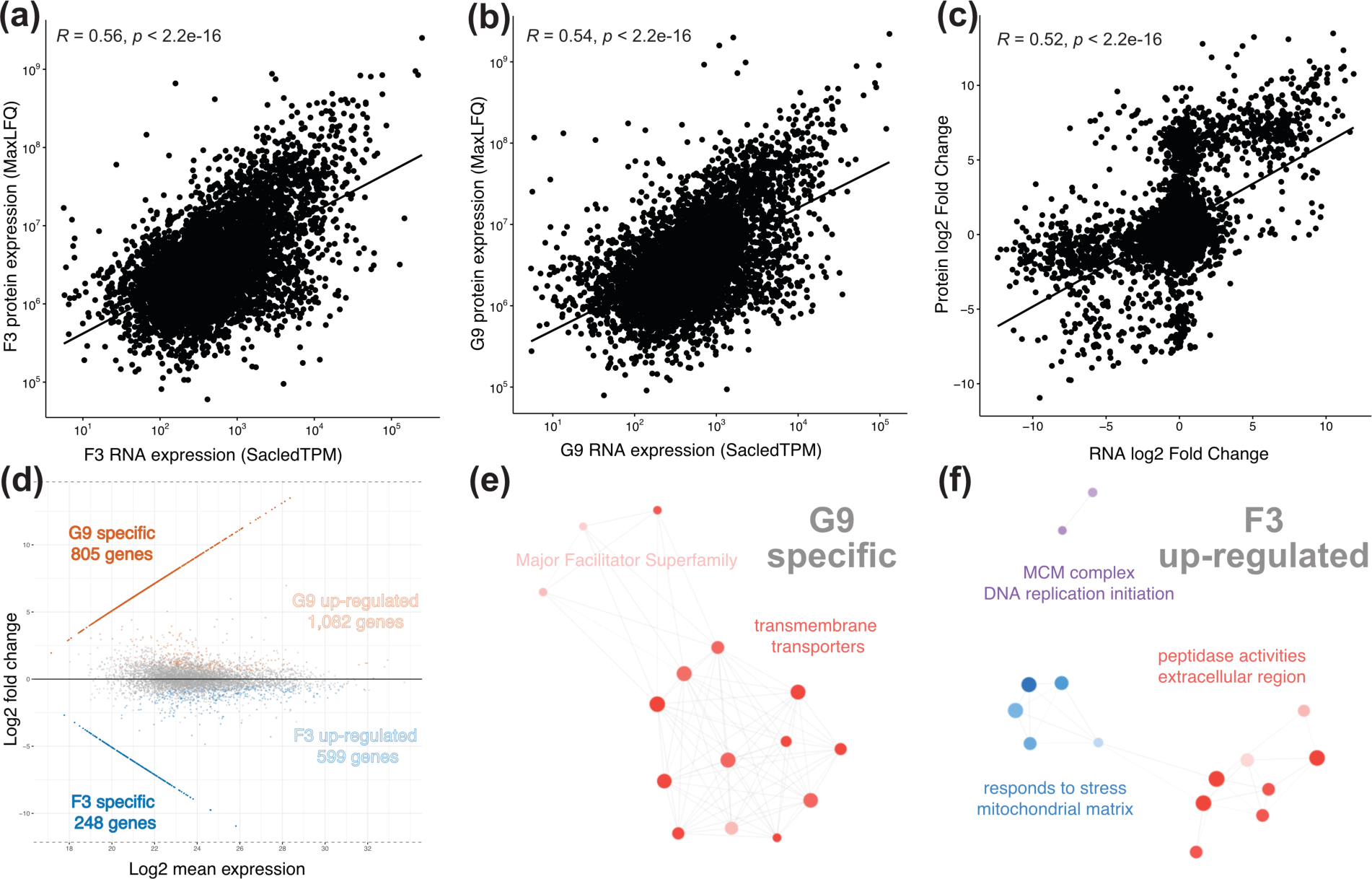
Protein-RNA correlations and proteomic differentiation. (a, b) Correlation between RNA and protein expression in F3 (a) and G9 (b). (c) Correlation between RNA and protein log_2_ fold change between F3 and G9. (d) MA plot of protein expression, highlighting genes specifically expressed or up-regulated in each life cycle phases. (e) Clusters of enriched GO terms of diploid-specific proteins. Each dot represents an enriched term, with darker colors indicating smaller p-values. Dots with similarity > 0.5 are linked by a grey line. Clusters with ≤ 5 genes were excluded. (f) Clusters of enriched GO terms of up-regulated proteins in the haploid.

### DNA methylation profiles and gene expression

Whole-genome 5-methylcytosine (5mC) patterns were profiled using two replicates of bisulfite sequencing for the clonal populations of F3 and G9 (Table S1). DNA methylation is prevalent in symmetric contexts, with CG and CHG sites accounting for 95% of the methylated sites (Fig. 5a). In the haploid, almost all sites are either lowly methylated (methylated reads < 20%, accounting for 80% of CG and 90% of CHG) or highly methylated (methylated reads > 80%, accounting for 18% of CG and 9% of CHG). The diploid has lowly methylated (76% of CG and 89% of CHG) and highly methylated (12% of CG and 5% of CHG) sites, along with sites displaying intermediate methylation levels (methylated reads between 20% to 80%; 11% of CG and 5% of CHG) (Fig. 5b). Cross-referencing with the genomic variation data indicates only a negligible number of intermediate-methylated sites resulting from transitions in one of the two alleles (2,090 T-C and 2,000 A-G transition sites corresponding to 0.2% of all intermediate-methylated sites). Instead, they are mainly caused by differential methylation on sister chromatids.

**Fig. 5.**
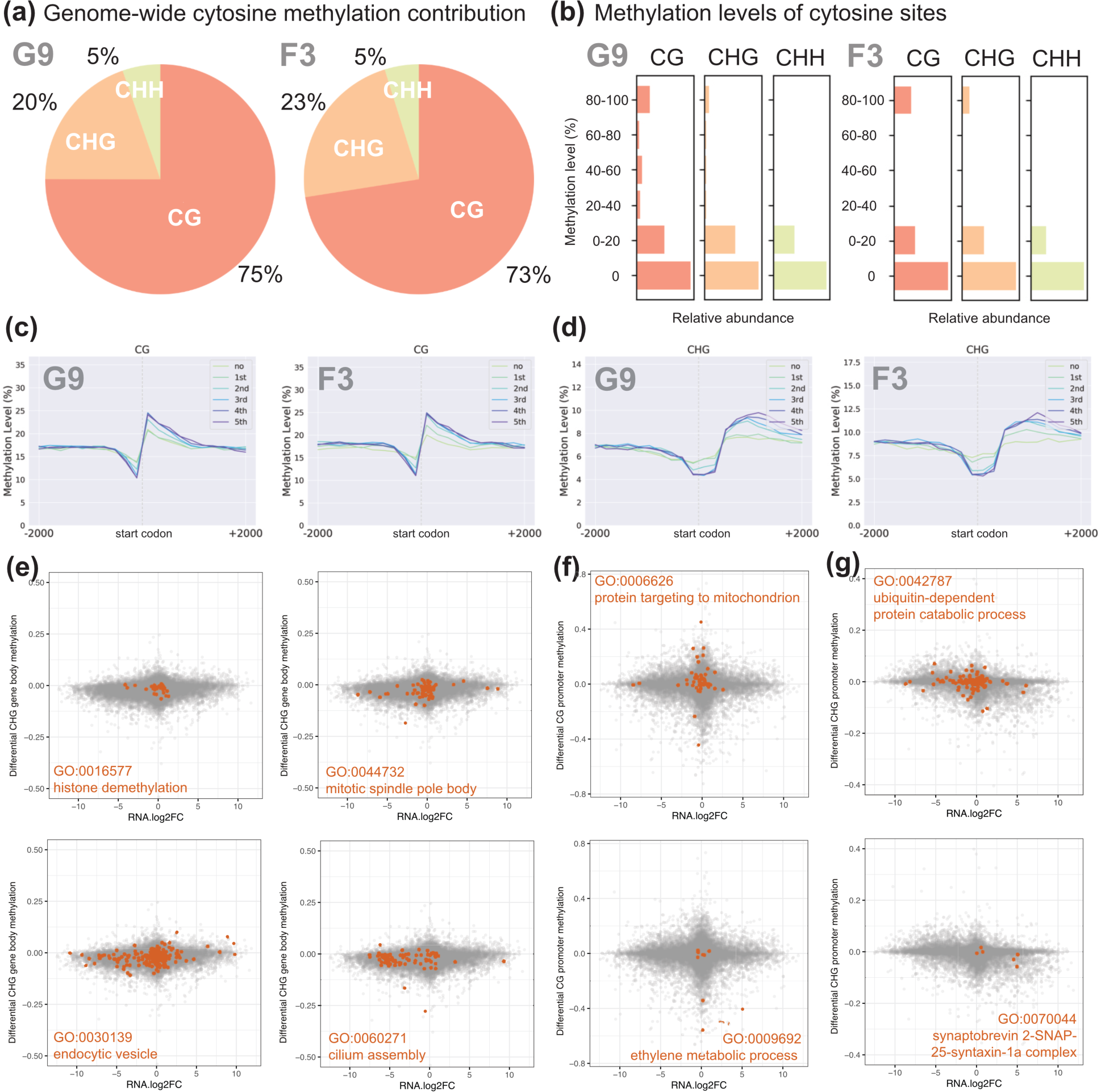
Genome-wide methylcytosine (mC) patterns, levels, and correlations with RNA expression. (a) Percentage of mC contribution in different sequence contexts: CG, CHG, and CHH. (b) Relative abundance of sites with different mC levels in CG, CHG, and CHH sequence contexts. (c, d) Average mC level profiles (CG [c] or CHG [d] context) around genes in six categories of RNA expression (5^th^: highest level). (e) Selected enriched GO terms among genes with higher RNA and higher gene-body mC levels in one life cycle phase (CHG context). (f, g) Selected enriched GO terms among genes with higher RNA and lower promoter-region mC levels in one life cycle phase (CG [f] or CHG [g] context).

DNA methylation of transposable elements (TEs) has been linked to heterochromatin formation and can influence life cycle phase differentiation (Vigneau & Borg, 2021). However, we did not observe a higher methylation level of TEs (Fig. S3). High CG methylation was observed in some rRNA genes, but not in other repetitive sequences (Fig. S3). Of the 43 genes annotated with DNA methyltransferase activity (GO:009008), two are up-regulated in the diploid and seven in the haploid. Notably, all three N-6 adenine methyltransferases (EHUX_g4335, EHUX_g18645, EHUX_g32175) are up-regulated in the haploid. However, only a few reads showed N-6 adenine methylation signals in our PacBio HiFi reads without any specific pattern.

To analyze the correlation between DNA methylation and RNA expression, we divided genes into six groups, including unexpressed genes (“no”) and five bins of expressed genes ranked by scaled transcripts per million (scaledTPM) (Soneson *et al*., 2015) (low to high: groups 1 to 5). Groups with higher RNA expression are associated with lower mean CG site methylation levels upstream of the start codon and higher levels downstream (Fig. 5c). For CHG sites, higher RNA expression is linked to lower methylation levels around the start codon and higher levels in the gene body (Fig. 5d). In both life cycle phases, E3 ubiquitin-protein ligase is among the top 10% of genes with the highest mean gene body CHG methylation levels (Dataset S3). A comparison between the two life cycle phases reveals positive association between RNA expression and gene body CG or CHG methylation for specific functions (Dataset S3). For instance, genes related to histone demethylation, mitosis, endocytosis, and flagella assembly often exhibit higher RNA expression and higher methylation levels in the gene body of the haploid (Fig. 5e). Genes of certain functions with significant differences in promoter methylation levels show differential RNA expression in the opposite direction. For instance, mitochondrion-targeting and ubiquitin-dependent proteins tend to show higher RNA expression and lower methylation levels in the haploid, whereas ethylene metabolic and SNARE proteins tend to exhibit higher RNA expression and lower methylation levels in the diploid (Figs. 5f, 5g).

### Genome-level allelic imbalance of RNA expression in diploid

Allelic imbalance is the phenomenon where the two alleles of a gene in a diploid cell have different RNA expression levels (Wu *et al*., 2023). Among the 10,762 heterozygous genes, 1,809 genes (16.81%) show significant signals of imbalanced RNA expression (Fig. 6a). Through comparison with the genome sequence of the haploid strain F3, we designated one G9 allele as the F3-like allele and the other as the alternative allele. Among the genes with allelic imbalance, a slightly higher proportion of F3-like alleles are up-regulated compared with the alternative alleles (Fig. 6b). Compared with genes with F3-like alleles up-regulated (23%), a much high proportion (52%) of genes with up-regulated alternative alleles also higher expression in the diploid than the haploid (Fig. 6c). This observation suggests that a fraction (approximately 400 genes) of the differentially expressed genes identified in this study may be attributed to allelic imbalanced RNA expression.

**Fig. 6.**
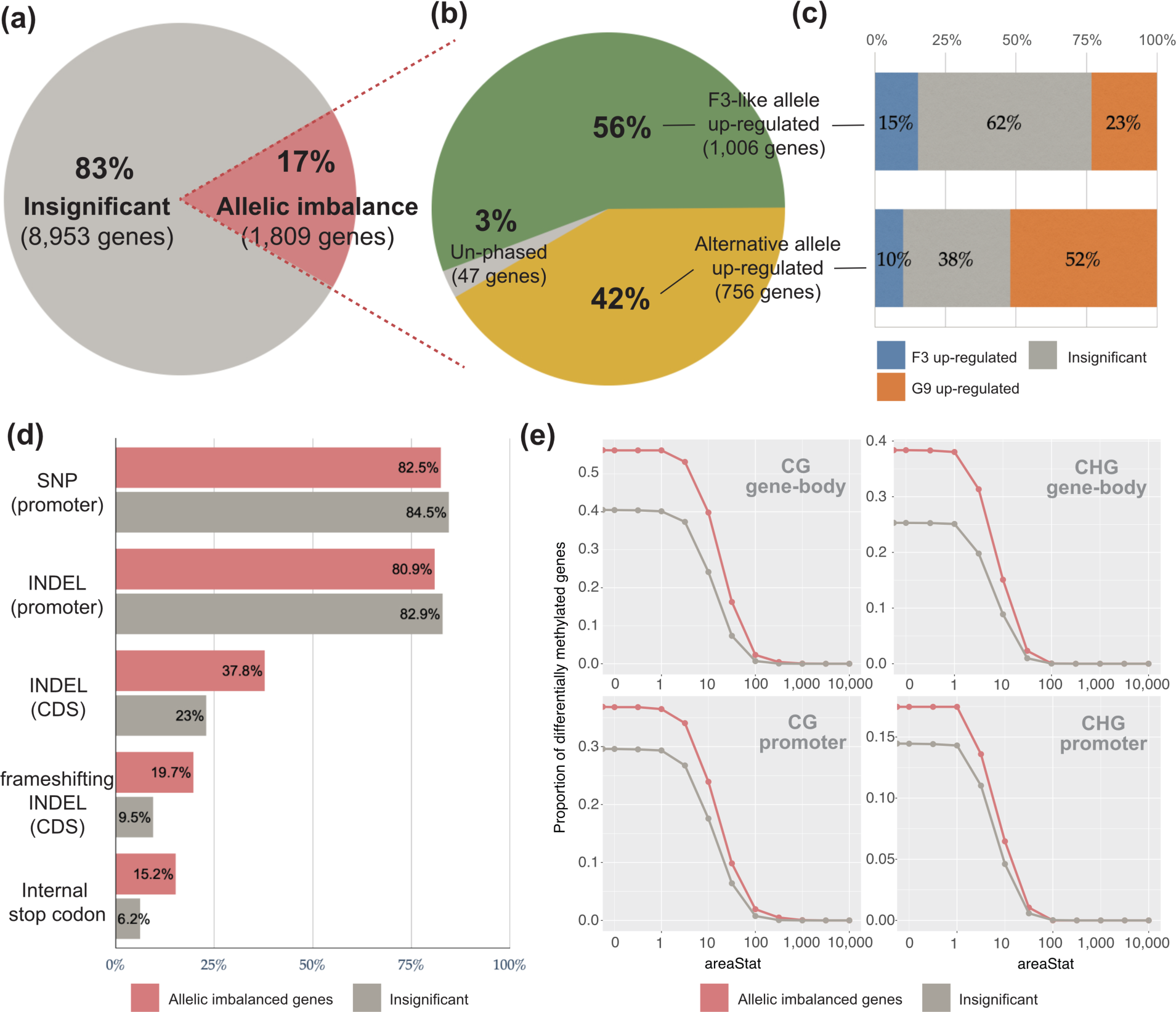
Allelic imbalance in RNA expression and its relationships with genomic variation and allele-specific methylation. (a) Percentage of allelic imbalanced genes among the heterozygous genes with exactly two copies in the G9 genome and with detectable transcripts. (b) Percentage of genes with up-regulated F3-like or alternative alleles among the allelic imbalanced genes. (c) The percentage of F3 up-regulated genes and G9 up-regulated genes among the genes with up-regulated F3-like or alternative alleles. (d) Percentage of genes with and without significant allelic imbalanced expression that contain single-nucleotide-polymorphisms (SNPs), insertion-deletions (INDELs), frameshifts, or internal stop codons. (e) Proportion of genes with and without significant allelic imbalance that have allele-specific methylated regions. AreaStat: the sum of t-statistics for the sites within the region.

The majority of the heterozygous genes have inter-allelic SNPs and/or indels (>80%) within 2,000-bp upstream of the start codon, but the percentage with allelic imbalance is not higher than without imbalance (Fig. 6d). However, we see a higher proportion of allelic imbalanced genes have indels in their coding sequences, particularly when these indels result in frameshifts and premature stop codons (Fig. 6d). We further examined the links between allelic imbalance and allele-specific methylation. A higher percentage of genes with allele-specific methylation patterns (in either the promoter region or gene body) was observed in genes with allelic imbalance than without in both CG and CHG contexts (Fig. 6e). Leveraging PacBio long-read DNA and RNA sequencing, we were able to better detect, phase, and visualize allelic imbalance and allele-specific methylation in a haplotype-resolved manner. For example, EHUX_g5658, a sulfatase gene with transcripts only detected in G9, displays allele-specific methylation in both the promoter region and gene body. Its F3-like allele exhibits the same methylation pattern as in the F3, and similar to the F3 gene, this allele is not transcribed (Fig. 7).

**Fig. 7.**
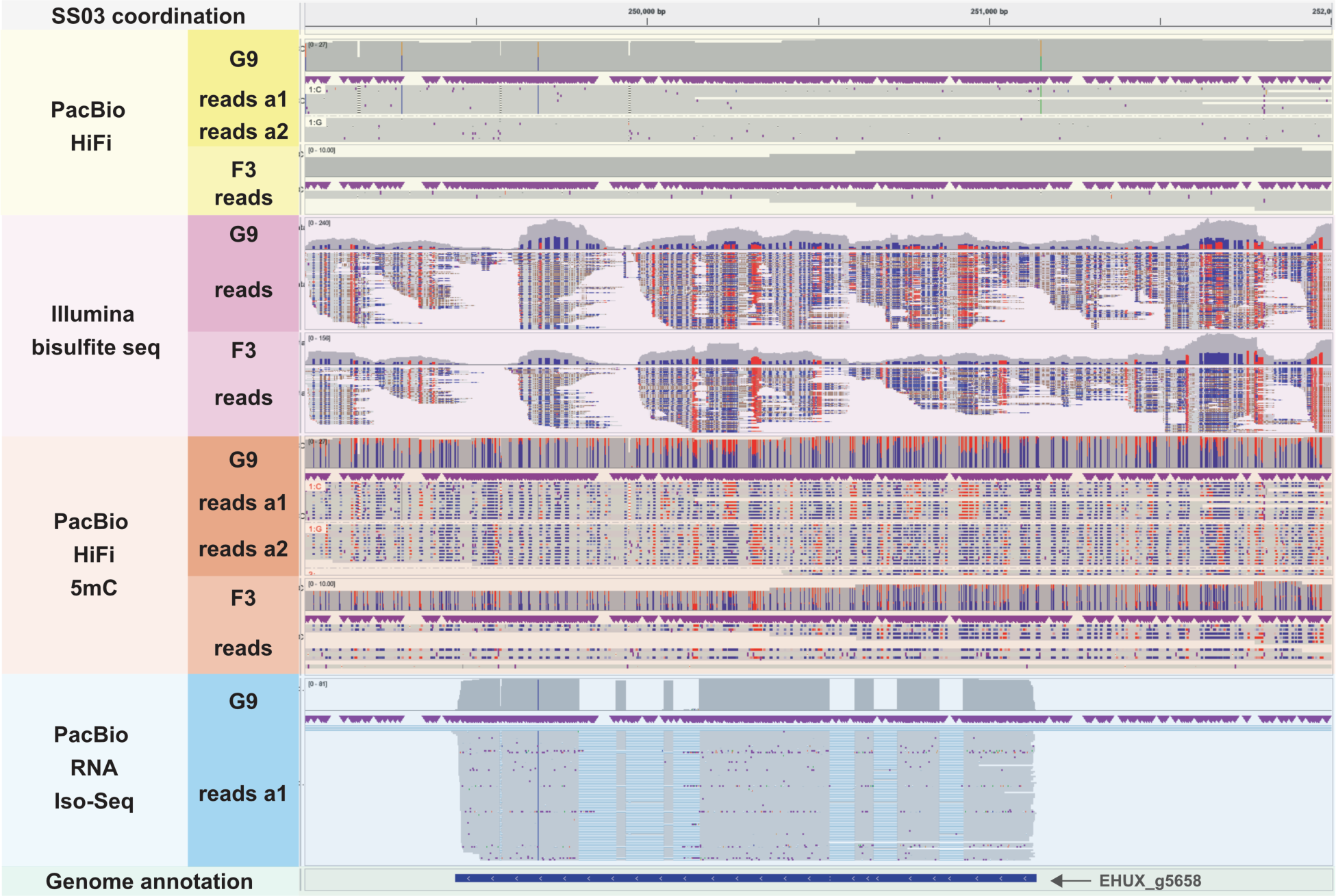
Allele-specific methylation and allele-imbalanced RNA expression in a *G*. *huxleyi* gene. Genomic, methylomic, and transcriptomic reads were mapped onto SS03 (249,000–252,000) containing EHUX_g5658, a sulfatase gene located on the Crick strand. Track yellow (top): PacBio HiFi reads (variable sites: A [green], C [blue], G [yellow], T [red], insertions [purple vertical bars], and deletions [black dashes]). Track pink: Illumina bisulfite sequencing reads (5mC [red], C [blue], or antisense [methylation status unknown; grey]). Track orange: PacBio HiFi reads with methylation calling (5mC probability > 50% [red] or 5mC probability < 50% [blue]). Track blue: PacBio Iso-Seq reads (transcript [grey] or intron [blue]). Track green: gene annotation.

### Histones and histone modifications

Epigenetic modifications of histones have been proposed as key players in life cycle phase transitions (Vigneau & Borg, 2021). Multiple copies of each core histone protein, including H2A, H2B, H3, and H4, were identified, but the H1 linker protein, which is found in animals, fungi, plants, and kinetoplastids (Discoba) (Talbert *et al*., 2019), is not present in the *G*. *huxleyi* or *G*. *pseudohuxleyi* genomes. Both haploid and diploid phases exhibited up-regulation of specific histone paralogs, as well as histone lysine methyltransferases and deacetylases (Dataset S4). Genes associated with H3K9 methylation are known to influence life cycle phase differentiation in plants, green algae, and diatoms (Vigneau & Borg, 2021). In our study, the transcripts of K9 histone-lysine N-methyltransferase SUV39H2 and the protein of enhancer of zeste E(z) were exclusively expressed in the diploid phase. Two other K9 histone-lysine methyltransferases (EHUX_g17408 and EHUX_g28648) and some histone arginine N-methyltransferases (PRMTs) are also up-regulated in the diploid.

### Diploid-haploid physiological differentiation

We summarized the multiomic profiles of the diploid and haploid phase of *G*. *huxleyi* by focusing on genes related to various physiological functions (Fig. 8; Notes S1, S2, and Dataset S4) (Mackinder *et al*., 2011; Benner *et al*., 2013; Taylor *et al*., 2017; Brownlee *et al*., 2021; Skeffington *et al*., 2023). The diploid and haploid show clear differences in certain carbonic anhydrases, transporters, and SNAP receptors (SNAREs) that are involved in coccolithogenesis. We also demonstrate that they utilize different sets of genes involved in polysaccharide and sulfate metabolism. The considerably higher concentration of DMSP in the diploid (Heidenreich *et al*., 2019) is attributable to more genes in DMSP degradation pathways being upregulated in the haploid. In line with a more mixotrophic nutrition, genes related to “endocytic vesicle” (GO:0030139) are up-regulated and have lower CHG methylation in the haploid. On the contrary, many photosynthesis-related genes, including those for chlorophyll synthesis and light harvesting, are up-regulated in the diploid.

**Fig. 8.**
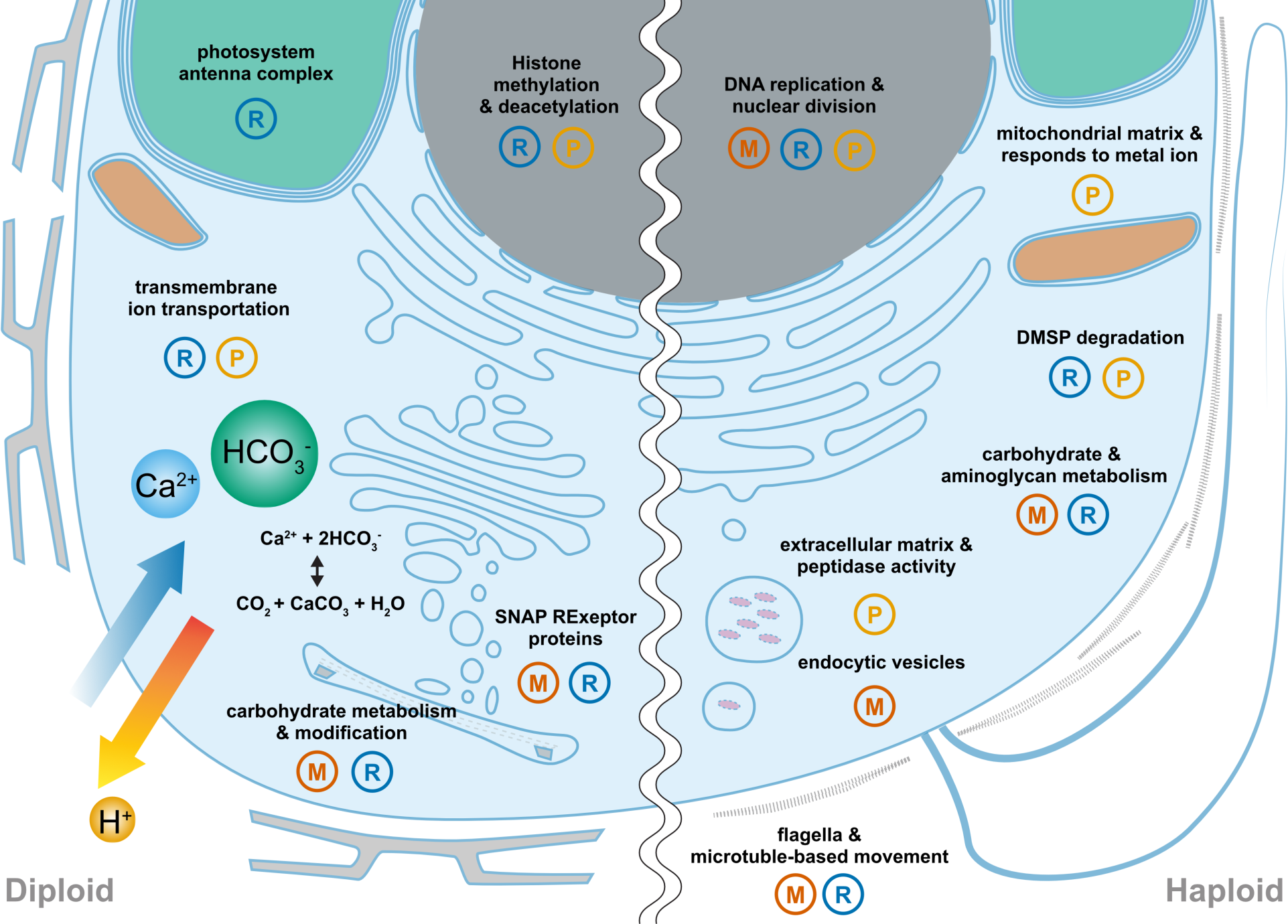
Summary of molecular differentiation between the diploid (left) and haploid (right) cell. The types of data supporting each functional category are indicated. M: DNA cytosine methylation; R: mRNA expression; P: protein expression.

## Discussion

Our multiomic sequencing and analyses lay a robust foundation for advancing coccolithophore research. Highly contiguous *de nov*o genome assemblies serve a critical role in dissecting the diploid-haploid variation in the life cycle of *G*. *huxleyi*. Profiling and comparing the transcriptomes, proteomes, and methylomes further provide insights into potential gene expression regulatory mechanisms behind morphological, physiological, and ecological differentiation of diploids and haploids. This study underscores the importance of genome phasing in comparative genomics and multiomics, and carries profound implications for future investigations into haploid-diploid differentiation and calcification mechanisms in coccolithophores.

Consistent with known high variation in genome size, structure, and gene content within *Gephyrocarpsa* and haptophytes (Read *et al*., 2013; Yang *et al*., 2023), more than 3,000 (13%) orthologous gene group identified in *G. huxleyi* are absent in *G. pseudohuxleyi* (Fig. 2c). The genomes of RCC1216 and RCC1217, a haploid-diploid pair separated by approximately twenty years, unveiled notable copy number differences at the superscaffold level and hundreds of structural variations, including two instances of chromosome-level recombination (Fig. 2e; Fig. S1). These findings highlight the prevalence of structural variation between closely related coccolithophore strains (Note S3). Instead of meiosis and sexual reproduction, the high frequency of variations is more likely due to homologous recombinations during DNA repair (mitotic recombinations), as has been shown in diatoms (33). Still, the Genomic copy numbers and structural variations (Fig 2e, 2f; Dateset S1) can be potential post-zygotic barriers to nuclear gene flow, offering a plausible explanation for the observed restricted nuclear gene flow among morphotypes within *Gephyrocapsa* (Filatov *et al*., 2021; Kao *et al*., 2022). In line with DNA content estimation based on flow cytometry (Houdan *et al*., 2004; Frada *et al*., 2017), we show a higher frequency of insertions in F3 (or deletions in G9) than F3 deletions (or G9 insertions) (Dateset S1), which leads to a diploid genome size smaller than twice the haploid genome size (Table 1).

The diploid-haploid differentiation in *G. huxleyi* involves genes encoding diverse cellular and biochemical functions (Fig. 8; Notes S1, S2, Dataset S4). The enrichment of transmembrane transporters among up-regulated diploid genes underscores the crucial role of ion transporters in calcification, including bicarbonate transporters (SLC4, ABCC8), calcium ion transporters and channels (CAX3, SLC24, TRPM), and H^+^/Na^+^ exchangers and a vacuolar-type ATPase for removal of the byproduct proton to maintain pH homeostasis. For the exocytosis of coccoliths, SNAP receptors (SNAREs) that mediate vesicle fusion with target membranes show diploid-specific expression (Fig. 8; Dataset S4) (von Dassow *et al*., 2009). We identified 15 carbonic anhydrases (CAs), which are pivotal for the conversion between carbon dioxide and bicarbonate and potentially regulate carbon acquisition and allocation between photosynthesis and calcification. Among the four CA families, αCAs and δCAs exhibit up-regulation in the diploid phase and are likely to be directly involved in calcification.

Regulation of CaCO_3_ crystal growth on an organic baseplate within the coccolith vesicle involve both proteins and polysaccharides. As the major organic component of coccoliths, coccolith-associated polysaccharides (CAPs) are acidic, can contain sulfate groups, and mediate the calcification process by directing calcium ions and controlling crystal morphology (Marsh *et al*., 2002; Gal *et al*., 2016; Marzec *et al*., 2019). Transcriptomic differences between the diploid and haploid strains are marked by an enrichment of enzymes for carbohydrate metabolism (including glycosyltransferases), sulfatases, and sulfotransferases (Fig. 3; Dataset S3), suggesting that diploid and haploid cells have a major difference in their carbohydrate components that might be related to coccolithogenesis. The diploid-haploid differential expression can be regulated through allele-imbalanced transcription (Fig. 6), which is in turn associated with allele-specific methylation patterns in the diploid (Fig. 7). To further elucidate the mechanisms for epigenomic reprograming in *G*. *huxleyi*, it remains a major challenge to systematically manipulate diploid-haploid transitions in the lab.

In addition to coccolithogenesis, the transcriptomic and proteomic profiles reflect other physiological differences between the haploid and diploid phases (Fig. 8; Note S2; Dataset S4). The presence of flagella and a likely more mixotrophic nutrition in the haploid are in contrast with more abundant chlorophyll a under high irradiance (Hariskos et al., 2015; Ruan et al., 2023; 300 and 100 μmol photons m^−2^ s^−1^, respectively) and the up-regulation of photosynthesis-related genes in the diploid. The low abundance of DMSP in the haploid (Heidenreich *et al*., 2019) is possibly due to the up-regulation of DMSP catabolic enzymes in both demethylation and cleavage pathways, including two key DMSP lyases, DddD and Alma. Given that DMSP and derived compounds are known to mediate microbial interactions (Alcolombri *et al*., 2015; Barak-Gavish *et al*., 2018), differences in DMSP metabolism imply that the two life-cycle phases might differ in their microalga-microbe interactions, as also suggested by the different bacterial composition associated with G9 and F3 cultures (Table S2). Consistent with the previous observation that haploids show a higher growth rate under higher irradiance (Hariskos *et al*., 2015), we found that MCM, crucial for DNA replication initiation, is enriched among the haploid up-regulated proteins (Fig. 4f; Dataset S4), suggesting increased mitotic activity in F3. Together these physiological differences point to potential ecological differentiation between the diploid and haploid, which merits further investigation using field-collected data.

Epigenetic modifications of DNA and histones have been proposed as key players in life cycle phase transitions (Vigneau & Borg, 2021), with varying patterns across the tree of eukaryotes. Animals typically exhibit methylation in the CG context, while land plants and the green alga *Chlamydomonas* manifest methylation in CG, CHG, and CHH contexts (Kawashima & Berger, 2014; Lopez *et al*., 2015). In contrast to methylation in red algae (bias towards non-CG) (Lee *et al*., 2018), diatoms (predominantly CG) (Veluchamy *et al*., 2013; Traller *et al*., 2016), and brown algae (absent or minimal and confined to CHH) (Cock *et al*., 2010; Fan *et al*., 2020). *G. huxleyi* methylation is found in CG (∼18%) and CHG (∼9%) contexts, with relatively rare CHH methylation (∼1%) (Figs. 5a, 5b, S3). We show that methylation patterns can be heritable, as evidenced by the similarity between the F3-like haplotype in G9 and the F3 genome in certain regions after at least 20 years of separation (Fig. 7). Unlike Chloroplastida, where gene body CG methylation up-regulates gene expression and methylation in CHG contexts is typically associated with low expression (Bewick & Schmitz, 2017), genes with higher expression levels in *G. huxleyi* display increased methylation in both CG and CHG contexts (Fig. 5c, 5d). Histones H2A, H2B, H3, and H4 were identified within the genome, and each core histone class has specific paralogs up-regulated in either haploid or diploid cells (Dataset S4). In green algae and bryophytes, the regulation of life cycle phase transition has been suggested to involve histone modifications, particularly the tri-methylation of lysine 27 on histone H3 protein (H3K27me3) that silences gene expression (Vigneau & Borg, 2021). The up-regulation of H3K9 methyltransferase SUV39H2, enhancer of zeste E(z), and arginine methyltransferases in the diploid of *G*. *huxleyi* implies a potential role of histone methylation in diploid-haploid differentiation, but further studies are needed to profile histone modifications and test this hypothesis.

This study presents a comprehensive view of the life-cycle differentiation between diploid and haploid phases of *Gephyrocapsa* — globally distributed and highly abundant calcifying algae with pivotal roles in marine carbon cycling. In particular, our results show that (1) their genomes are dynamic and prone to variations, including chromosome-level structural variations, biased indel distribution between the diploid and haploid, and loss of heterozygosity in the diploid, as revealed by their highly contiguous *de novo* genome assemblies; (2) Transcriptomic and proteomic differences of diploid and haploid cells modulate their differentiation in coccolithogenesis, flagellar assembly and movement, photosynthesis, carbohydrate metabolism, DMSP degradation, DNA replication, and endomembrane system and transport; (3) Gene transcript abundance is correlated with patterns of DNA methylation, which occurs in CG and CHG contexts and can be inheritable, allele-specific, and differentiated between life-cycle phases. Haplodiplontic life cycles are widespread across diverse lineages of eukaryotes, where coccolithophores exemplify both algae and microbial eukaryotes that exist in two morphologically and physiologically distinct ploidy phases. By integrating genomic, transcriptomic, proteomic, and methylomic analyses, this study not only illuminates the molecular basis of diploid-haploid differentiation and maintenance in coccolithophores, but lays the groundwork for future research on their life cycling in marine ecosystems and its implications for the distribution, ecological roles, and microbial interactions of coccolithophores.

## Acknowledgments

We thank Pui-Yan Kwok, Hsiao-Jung Kao, and Hsiao-Huei Chen for their generous help with the Bionano optical mapping, Irina V. Agarkova and James L. Van Etten for advice on HMW DNA extraction, Chih-Horng Kuo for sharing their computing equipment, and Assaf Vardi for sharing the cultures of RCC1217 and RCC1216. We are also grateful for the sequencing services and technical support provided by BRCAS NGS High Throughput Genomic Core, IPMB Genomic Technology Core, National Center for Genomic Medicines, and Biotools. We acknowledge Yu-Fang Tseng of IPMB Cell Biology Core for flow cytometry analysis and sorting services, Wann-Neng Jane and the staff of the EM division for EM sample preparation, Chuan-Chih Hsu and IPMB Proteomics Core for their assistance with global proteomic quantification, Academia Sinica Common Mass Spectrometry Facilities for proteomics and protein modification analysis facilities, and the National Center for High-performance Computing and IPMB Bioinformatics Core Lab for computer time and facilities. We appreciate the consultation on genome-wide methylation analyses by the lab of Pao-Yang Chen, advice on genome assembling strategies by the lab of Chung-Yen Lin, and the suggestions of Michael Borg on the manuscript,. Lastly, we thank the help of other lab members with this project, especially Bo-Wei Wang and Elias Miguel Barreira. This work was supported by the intramural funding of the Institute of Plant and Microbial Biology (CK), Academia Sinica Career Development Award (AS-CDA-110-L01 to CK), Academia Sinica Postdoctoral Scholar Program (AS-PD-11001-L007 and AS-PD-11201-L01 to T.K.), and National Science and Technology Council, Taiwan (108-2311-B-001-040-MY3 and 111-2611-M001-008-MY3 to C.K.). The funding bodies had no role in the design of the study, in the collection, analysis, and interpretation of data, or in writing the manuscript.

## Author contributions

This project was initiated and conceptualized by CK and MF. The primary research activities, including DNA extraction, sequencing, genome assembly, data analysis, and synthesis were conducted by TK. ML was responsible for RNA purification for Illumina RNA sequencing and PacBio Iso-Seq, as well as preparing cell cultures for TEM observation and global protein identification. TW contributed to maintaining cell cultures, participated in genome assembly and annotation, and assisted with data curation. CL played a key role in the establishment of the HMW DNA extraction protocol. The initial draft of the manuscript was prepared by CK and TK. All authors contributed their insights, reviewed the manuscript, and provided valuable input, ultimately approving the final version. Throughout the project, CK provided supervision and guidance.

## Data availability

All data are available in the main text or the supplementary materials. The genome assemblies have been deposited in the NCBI under BioProject PRJNA774110 and the accession numbers are listed in Table S1.

